# Intravenous psilocybin administration attenuates mechanical hypersensitivity in a rat model of chronic pain

**DOI:** 10.1101/2023.08.26.554802

**Authors:** Nicholas Kolbman, Tiecheng Liu, Peter Guzzo, Jim P. Gilligan, George A. Mashour, Giancarlo Vanini, Dinesh Pal

## Abstract

There is a renewed interest in the therapeutic potential of psychedelics, including psilocybin, in treating mental health disorders. However, there are no data on the efficacy of psilocybin in alleviating chronic pain. In this study, we investigated the effect of psilocybin on mechanical hypersensitivity and thermal hyperalgesia in a rat model of formalin-induced chronic pain. Adult male and female rats were surgically implanted with a jugular vein catheter for psilocybin or saline administration. After two weeks of post-surgical recovery and conditioning, baseline responses to mechanical (von Frey assay) and thermal (hot plate assay) stimuli were measured. Twenty-four hours after baseline measurements, rats received a subcutaneous injection of formalin (5%, 50µL) into one of the hind paws and 2h later, responses to the mechanical and thermal stimuli were measured. Twenty-four hours after formalin injection, rats received an intravenous bolus of 1 mg/kg psilocybin (n=14) or 10 mg/kg psilocybin (n=12) or saline (n=13), and approximately 3h later, responses to the mechanical and thermal stimuli were measured. Rats were tested every other day during week 1, and then weekly for the next 3 weeks. Formalin injection induced thermal hyperalgesia and bilateral mechanical hypersensitivity in the hind paws of all rats. Intravenous psilocybin produced significant attenuation (p<0.05) of the formalin-induced bilateral mechanical hypersensitivity for 28 days but had limited effect (p<0.05 only on days 1, 3, 5, and 21) on thermal hyperalgesia. These data demonstrate that a single intravenous bolus of psilocybin can attenuate indices of chronic pain in a rat model.

---

There is a renewed interest in psychedelic drugs as potential therapeutic agents for the treatment of psychiatric disorders.^1^ In particular, psilocybin has shown promise for the treatment of refractory depression^2^ and major depressive disorder,^3^ and has also been explored as a treatment for tobacco and alcohol abuse.^4,5^ However, despite anecdotal reports,^6^ there has been no systematic study to investigate the effectiveness of psychedelics in chronic pain, a major public health problem. To address this gap, we determined the effect of intravenous psilocybin administration (low dose: 1 mg/kg, high dose: 10 mg/kg) on mechanical hypersensitivity and thermal hyperalgesia in a well-established rat model of formalin-induced, centralized chronic pain.^7,8^ In this rat model, formalin injection in one of the hind paws produces hypersensitivity in both the injected and non-injected paw, which was reported to persist for at least 28 days.^7,8^

The experimental design is illustrated in **Figure 1A**, and the detailed methodology is provided in the supplemental section. In brief, we first measured baseline responses to mechanical and thermal stimuli (adult rats, n=42, male=21, female=21) using the von Frey assay and hot plate assay, respectively. The response to mechanical stimuli – evoked by application of von Frey filaments of increasing thickness to the plantar surface of the hind paws – was quantified as paw withdrawal threshold. The response to thermal stimulus was quantified as the latency to paw withdrawal after being placed on a hot surface (52.5°C). Twenty-four hours after baseline measurements, all rats received a subcutaneous injection of formalin (5%, 50µL) into the dorsum of one of the hind paws (Day 0), and 2h after the formalin injection (the same day) we measured the responses to the mechanical and thermal stimuli. Thereafter, the rats were divided into three groups and 24h later (Day 1), **Group 1** (n=14, 6 male, 8 female) received an intravenous bolus of 1 mg/kg psilocybin, **Group 2** (n=12, 6 male, 6 female) received an intravenous bolus of 10 mg/kg psilocybin, and **Group 3** (n=13, 6 male, 7 female) received an intravenous bolus of 0.9% saline (vehicle control). Approximately 3h after psilocybin or vehicle infusion, we measured responses to mechanical and thermal stimuli (Day 1); thereafter the responses were measured every other day during week 1, and then weekly for the next three weeks.

**Figure 1:**
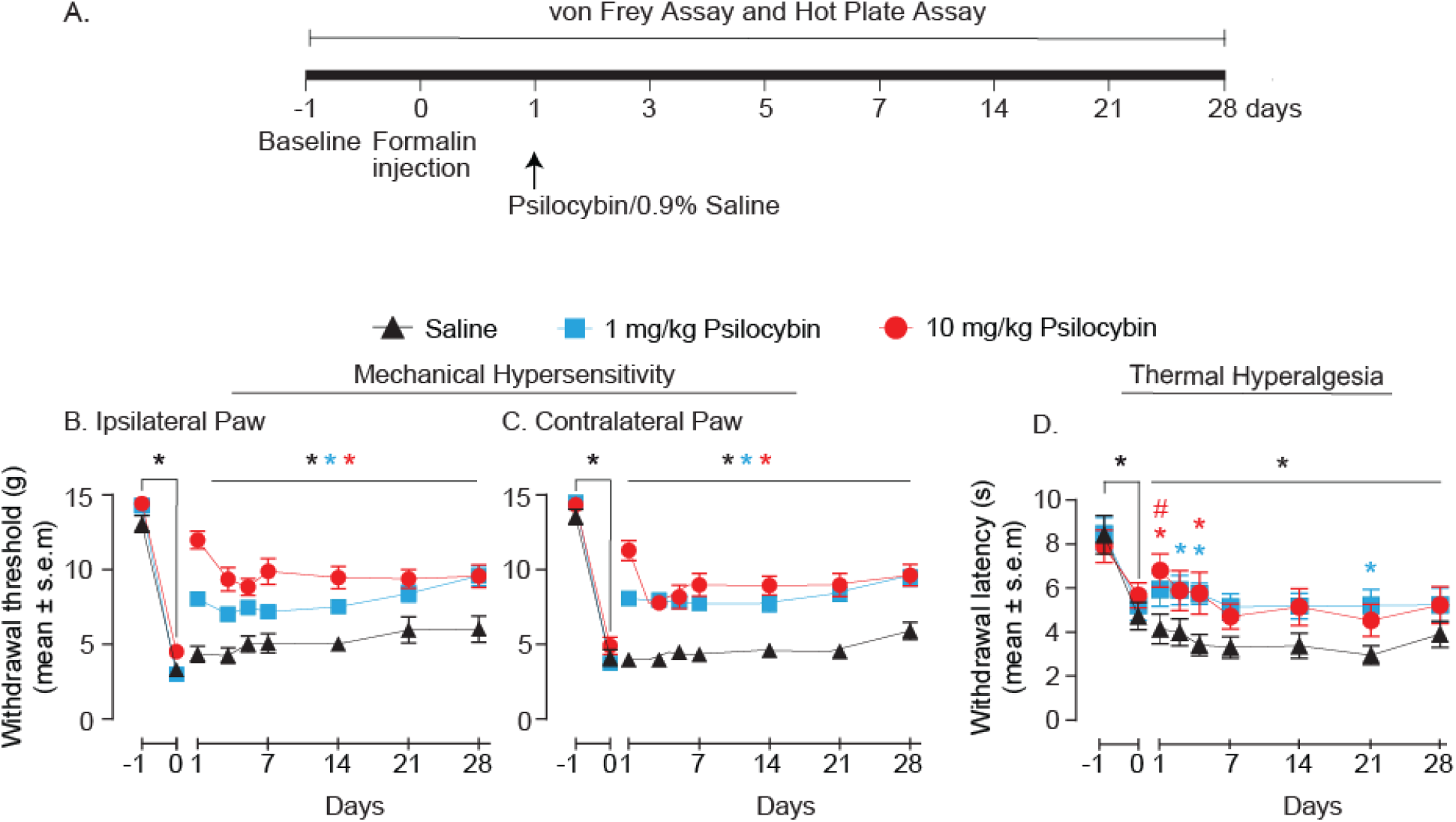
Schematic illustrating the design and timeline of the experiments **(A)**. Intravenous psilocybin attenuates formalin-induced bilateral mechanical hypersensitivity **(B-C)**, but not thermal hyperalgesia **(D)** in rat. As compared to the rats that received intravenous saline (black triangles, n=13, 6 male, 7 female), rats treated with the low (1 mg/kg – blue squares, n=14, 6 male, 8 female) or high (10 mg/kg – red circles, n=12, 6 male, 6 female) dose of psilocybin showed a significant attenuation of the formalin-induced mechanical hypersensitivity in both hind paws, which persisted for 28 days after formalin injection **(B-C)**. Treatment with the high dose of psilocybin (n=12, 6 male, 6 female) significantly attenuated formalin-induced thermal hyperalgesia, but the effect was restricted only to the day of psilocybin administration and day 5 after formalin injection **(D)**. Rats treated with the low dose of psilocybin (n=14, 6 male, 8 female) displayed a significant attenuation in the formalin-induced thermal hyperalgesia only on day 3, day 5, and day 21 after formalin injection **(D)**. Asterisks are color-coded to match the symbol color used for each group and represents statistical significance at p < 0.05. The exact p values are provided in the results section.

Similar data trends were observed between male and female cohorts and therefore both sex groups were combined for the statistical analyses using a linear mixed model (see the Supplemental Content for details). Formalin injection in either of the hind paws produced central sensitization to mechanical stimuli as demonstrated by a significant decrease (p<0.0001) in withdrawal threshold in the hind paws ipsilateral as well as contralateral to the injection (**Fig. 1B-C**). Similarly, formalin injection in either of the hind paws decreased (p<0.01) paw withdrawal latency upon application of thermal stimulus (**Fig. 1D**). Of note, both formalin-induced mechanical hypersensitivity and thermal hyperalgesia were observed for 28 days, which aligns with previously published data.^7,8^ Compared to the rats that received vehicle, the rats treated with psilocybin – low or high dose – showed a significant attenuation of formalin-induced mechanical hypersensitivity in both ipsilateral (p<0.01, all testing days) and contralateral (p<0.0001, all testing days) paws (**Fig. 1B-C**). In contrast to the effect on mechanical hypersensitivity, psilocybin showed limited efficacy in attenuating formalin-induced thermal hyperalgesia. Compared to the rats that received vehicle, the rats treated with high-dose psilocybin showed a significant attenuation of formalin-induced thermal hyperalgesia only on the day of psilocybin administration (p=0.0082) and on day 5 after formalin injection (p=0.02) (**Fig. 1D**). Low-dose psilocybin attenuated formalin-induced thermal hyperalgesia only on day 3 (p=0.04), day 5 (p=0.01), and day 21 (p=0.01) post-formalin injection (**Fig. 1D**). Psilocybin administration in naïve rats (i.e., without any formalin injection; n=3 male) did not alter sensitivity to mechanical or thermal stimuli.

Our results demonstrate that a single intravenous administration of psilocybin can attenuate mechanical hypersensitivity in a rat model of formalin-induced chronic pain for 28 days. Attenuation of centralized hypersensitivity (i.e., ipsilateral and contralateral to the site of formalin injection) suggests that psilocybin could act via neuroplastic effects produced by psychedelics.^9^ This is supported by the observation that psilocybin-induced attenuation of mechanical hypersensitivity outlasted the half-life of psilocybin (∼2h post intravenous administration),^10^ which suggests a centrally mediated mechanism of action. This is of translational relevance because many chronic pain conditions are posited to be the result of centralized neuroplastic changes in the brain and spinal cord. It is also important to note that psilocybin alone does not appear to influence pain indices in naïve rats. As such, the antinociceptive effects observed in this study are unlikely to result from any direct effect of psilocybin adminstration on behavioral phenotype. By contrast, psilocybin administration did not attenuate formalin-induced persistent thermal hyperalgesia. Although it is possible that the effect of psilocybin is specific to nociceptive modality, a more parsimonious explanation could be that the thermal stimulus, set at 52.5°C in the hot plate assay, was supramaximal and precluded antinociceptive effects of psilocybin.

Our study has several caveats. First, we used a rat model that develops chronic pain due to chemically induced inflammatory processes. Although our findings provide proof-of-concept for the use of psychedelics to attenuate chronic pain, further studies are required using animal models with diverse root causes of chronic pain (e.g.,neuropathic pain, early life stress models). Second, although we did not see any divergence in the data from male and female rats, our study is likely not powered to perform a direct comparison between sexes. Third, measurement of cold allodynia could have provided insights into temperature sensitivity, which in this protocol might have been masked by high temperature in the hot plate assay.

In conclusion, to our knowledge this is the first preclinical study demonstrating the effectiveness of any psychedelic to attenuate indices of chronic pain. Further studies are warranted to assess other endpoints related to chronic pain in male and female subjects, and the molecular mechanisms of psilocybin’s therapeutic effect.

## Supporting information

Kolbman et al., 2023 Supplemental Content

## Author Contributions

N.K., G.V., and D.P. designed the research; N.K and T.L. performed the experiments; all authors contributed to drafting the article; N.K., G.A.M., G.V., and D.P. interpreted the data and wrote the article.

## Acknowledgements

We thank Dr. Chris Andrews (Consulting for Statistics, Computing & Analytics Research, University of Michigan, Ann Arbor, Michigan) for consultation and help with statistical analysis.

