## Supplementary material for "Intravenous psilocybin administration attenuates mechanical hypersensitivity in a rat model of chronic pain": Kolbman et al., 2023 Supplemental Content

**Materials and methods:**

All experimental procedures were approved by the Institutional Animal Care and Use Committee at the University of Michigan and were conducted in compliance with the Guide for the Care and Use of Laboratory Animals (National Academies Press, 8th Edition, Washington DC, 2011). The experiments were conducted on adult Sprague Dawley rats (300-350 g, Charles River Laboratories Inc., MA) of both sexes (male = 21, female = 21). The rats were housed in a temperature-controlled facility, provided with *ad libitum* food and water, and maintained on a 12 h:12 h light-dark cycle (lights on at: 08:00).

*Surgical procedures:*

The rats were placed in an air-tight clear rectangular chamber (10.0 inches × 4.8 inches × 4.2 inches) to induce general anesthesia with 4–5% isoflurane (Piramal Enterprises, Telangana, India) in 100% oxygen. The anesthetic concentration was titrated (1-2%) to maintain absence of pedal withdrawal reflex and was continuously monitored using an anesthetic agent analyzer (Datex Medical Instrumentation, Tewksbury, MA). Body temperature was monitored using a rectal probe (RET-2 ISO, Physitemp Instruments, Inc., Clifton, NJ) and maintained at 37.0 ± 1°C using a small animal far-infrared heating pad (Kent Scientific Co., Torrington, Connecticut). The rats were implanted with an indwelling catheter (Micro-Renathane tubing, MRE-040, Braintree Scientific, MA) in the internal jugular vein for intravenous infusion of 0.9% saline or psilocybin (Cayman Chemical, MI; CAS 520-52-5). The jugular venous catheter was flushed with 0.2 mL of heparinized (1 unit/mL, Sagent Pharmaceuticals, Schaumburg, IL) saline and locked with 0.05 mL of Taurolidine-Citrate Catheter lock solution (TCS-04, Access Technologies, Skokie, IL) every 5-7 days to maintain catheter patency. In addition, stainless-steel screw electrodes were implanted across frontal, parietal, and occipital cortices to acquire electroencephalogram (EEG) data. The EEG data were collected but were not analyzed or presented for the current study. The rats received carprofen (5 mg/kg, s.c., Hospira, Inc., Lake Forest, IL) and buprenorphine

(0.01 mg/kg, s.c., Buprenex, Reckitt Benckiser Pharmaceuticals, Richmond, VA) for pre-emptive presurgical analgesia, and cefazolin (West-Ward-Pharmaceutical, Eatontown, NJ) (20 mg/kg, s.c.) as a pre-surgical antibiotic. The rats received buprenorphine (0.03 mg/kg, s.c.) every 8–12 h for 48 h for post-surgical analgesia.

*Formalin administration:*

Formalin (10%, Thermo Fisher Scientific, Waltham, MA, USA; #SF100-4) was diluted to 5% in sterile normal saline and filtered with a 0.22-µm syringe filter (Fisher Scientific; #05-713-386) immediately prior to injection. Rats were gently restrained and administered a subcutaneous injection (50 µL, 26-gauge needle) of 5% formalin into the dorsal surface of one of the hind paws in a counterbalanced manner.

*Experimental design:*

The experimental design and timeline are illustrated in Figure 1. The rats were provided at least 7 days of post-surgical recovery period before undergoing conditioning to the von Frey apparatus and hot plate. The rats were conditioned to the von Frey testing apparatus for a minimum of 1 week prior to beginning experiments after which baseline responses (Day -1) to mechanical and thermal stimuli were assessed. Twenty-four hours later, rats received a subcutaneous injection of 5% formalin into the dorsal surface of one of the hind paws (Day 0). Two hours after the formalin injection, nociceptive responses to the mechanical and thermal stimuli were reassessed. Thereafter, the rats were divided into three groups and twenty-four hours later (Day 1), **Group 1** (n=14) received an intravenous bolus infusion of 1 mg/kg psilocybin, **Group 2** (n=12) received an intravenous bolus of 10 mg/kg psilocybin, and **Group 3** (n=13) received an intravenous bolus of 0.9% saline (vehicle). Approximately three hours after psilocybin or vehicle infusion, we measured responses to the mechanical and thermal stimuli (Day 1), and after that the responses were measured every other day during week 1, and then weekly for the next three weeks.

*Nociceptive testing:*

Nociceptive responses were obtained between 12:00 and 14:00 and were collected by an investigator blinded to the treatment condition. Mechanical sensitivity was always assessed prior to thermal hyperalgesia. For assessment of mechanical sensitivity, rats were allowed 20 to 30 minutes to habituate to von Frey testing apparatus before each experiment. Mechanical sensitivity was assessed using the ascending method and was expressed as paw withdrawal threshold in grams (g) of force, as described in detail in previous work from our group and others.[1-3] Briefly, rats were placed in individual Plexiglas testing enclosures on a IITC von Frey mesh stand (Life Science Inc., Woodland Hills, CA, USA). For each experiment, seven von Frey filaments (1, 2, 4, 6, 8, 10 and 15 g of force) were applied in ascending order to the plantar surface of the rat hind paw until bent (Touch Test Sensory Evaluator, North Coast Medical Inc., Gilroy, CA, USA). Filaments were applied to the central region of the plantar surface making sure to avoid the foot pads, and only when the rat was standing on all four paws. No stimuli were applied if the rat was walking, grooming, or sniffing. After the filament bent, it was held for 5 s, or until a withdrawal response occurred. Each filament was presented to each hind paw five times with a 30 s interval between each stimulus. A response was considered positive when the hind paw being assessed was completely removed from the testing platform. The mechanical threshold was defined as the average minimal amount of force required to evoke a paw withdrawal response on each paw after five trials. Thermal hyperalgesia was assessed with the hot plate assay.[4] A metal hot plate was heated at a constant temperature of 52.5 °C with a clear acrylic cylinder placed atop to confine the rats in a specified observation field. The hot plate and cylinder were wiped clean with water after each measurement. Thermal hyperalgesia was expressed as paw withdrawal latency in seconds, or specifically, time to a lick of the hind paw or overt escape attempt.

*Statistical analysis:*
All statistical analyses were conducted using R (software version 4.3.1, R Core Team, 2023) and in consultation with the Consulting for Statistics, Computing, and Analytics Research core at the University of Michigan (CSCAR). GraphPad Prism software version 9.5 (GraphPad Software, San Diego, CA, United States) was used to create all graphs. Data are reported as mean ± standard error of the mean. A linear mixed model fit with lme4 was used to compare the effect of psilocybin and vehicle on formalin-induced mechanical hypersensitivity and thermal hyperalgesia. The linear mixed model included dose (vehicle, 1 or 10 mg/kg psilocybin), measurements on different days (baseline: day -1, formalin injection: day 0, psilocybin/saline injection: day 1, and post-formalin testing days 3, 5, 7, 14, 21, and 28 post-formalin administration), and their interaction, as well as sex (male or female) and its interaction with day, all as fixed effects, and with the subject (rat) serving as a random intercept. Post-hoc tests were utilized to compare subjects treated with vehicle and either 1 or 10 mg/kg psilocybin on each day, averaged over sex. The linear mixed model was also used to determine whether subjects, on average, returned to baseline levels of mechanical hypersensitivity and thermal hyperalgesia by comparing subjects administered psilocybin or vehicle to their paired baseline measurements. All comparisons were considered statistically significant if $p < 0.05$.
